# Foraging associations are related with helping interactions in a cooperatively breeding bird

**DOI:** 10.1101/2024.10.28.620677

**Authors:** André C. Ferreira, Damien R. Farine, Liliana R. Silva, Rita Fortuna, Claire Doutrelant, Rita Covas

## Abstract

Kin selection has been the main hypothesis explaining helping behaviour in cooperative breeders, with evidence being largely based on the observation that helpers tend to provide to related offspring. However, kin-biased help could conceal additional, mechanisms contributing to the maintenance of cooperation. Under pay-to-stay, group augmentation and partner choice hypotheses, a range of direct benefits can arise through helping. Here, we explored this potential mechanism by testing whether the social associations of breeding individuals were related with the help that they received from non-breeding individuals. We collected social associations from PIT-tagged sociable weavers, *Philetairus socius*, at RFID feeding stations, which allowed us to compare associations between breeders and either their helpers (mostly kin) or their other kin that did not help—before, during and after reproduction. Using correlative tests and data-driven simulations, we show that helpers have stronger foraging bonds with breeders than non-helping kin, and that these stronger bonds are present both prior and post breeding. Furthermore, helper-breeding female social affiliations were positively correlated with the amount of help provided. Our results suggest that direct benefits of social associations complement kin selection to determine helping decision, and that these in turn influence future social associations.

## Background

Cooperation is expected to be evolutionarily stable if the advantages to co-operators outweigh the costs of the cooperative behaviours ([1]. These benefits are classified as direct or indirect, and are not mutually exclusive. Indirect benefits are payoffs derived from assisting genetically related individuals, thereby improving their fitness, and are central to Maynard Smith’s kin selection theory ([2]; inspired by Hamilton’s inclusive fitness theory [3]).

Direct benefits, on the other hand, encompass a variety of paybacks obtained from helping others that ultimately improve the (future) reproductive success or survival of the co-operator [4]. These benefits, however, can arise from a broader range of contexts and are challenging to measure [5]. For example, benefits can be derived from reciprocal interactions [6] in which the same cooperative actions are exchanged between individuals, such as the case of food sharing in vampire bats and humans (e.g. [7,8]) or allo-grooming in many primates (e.g. [9,10]). In addition, given that the social environment experienced by the individuals affects fitness [11–13], cooperation could also be beneficial if cooperative actions generate benefits arising from increasing the strength of social ties to recipients. Specifically, cooperating has been suggested to lead to the inclusion of the co-operator in social groups or territories, (the “pay-to-stay” hypothesis [14,15]), leading to increased chances of obtaining a sexual or social partner (“partner choice” hypothesis [16,17]), to influence the perception of others towards the co-operators’ reputation (“social prestige” hypothesis [18]), or to lead to an increase in social group size the “group augmentation” hypothesis [19].

In birds, one of the most commonly studied forms of cooperation is helping-at-the-nest. It is characterized by a social system in which non-breeding, sexually mature individuals (called “helpers”) contribute to rearing the offspring of other breeders [20,21]. Kin selection is largely accepted as the main explanation for the evolution of cooperative breeding in most bird species [21]. Different lines of empirical evidence supporting kin selection with the main one based on the observation that helpers tend to assist genetically related young, and that such help usually translates into the production of a larger number of genetically related individuals, a reduction in the genetically related breeders’ workload, and/or increased survival of the breeders [22]. Nevertheless, the contribution to increased survival of chicks and breeders is not only predicted by kin selection. Hypotheses relying on the benefits associated with social affiliations (“group augmentation”; “partner choice” and “pay-to-stay”) also predict that helpers will contribute to the survival and/or production of others thereby promoting the maintenance of the group or partnership in order to gain socially derived (direct) benefits. Yet, studies rarely consider that helping interactions among kin could arise not only because of kin associated benefits, but also as a consequence of the spatial and/or social organization of the population [23]. For example, social barriers to movements [24] limit opportunities to associate with, and therefore help, non-kin (e.g. “population viscosity” hypothesis; [3,25,26]). Furthermore, even if individuals have the choice between helping kin and non-kin, characteristics usually associated with relatedness, such as familiarity, tolerance towards related individuals or prior knowledge of the territory, could make relatives better social partners [27]. Thus, disentangling the effect of kinship from other social factors remains both a major hurdle and a major gap in our understanding of cooperation.

Direct benefits arising from social associations are inherently more challenging to study than indirect benefits, as collecting data on social ties requires many observations of interactions among individuals, whereas relatedness among all study individuals can be determined using just one sample per individual (see [5]). Consequently, current methods and statistical tools may bias results towards favouring kin selection, while the contribution of social associations in promoting the evolution or maintenance of cooperation is likely to have been systematically under-estimated. For example, Carter et al., [5] used simulated and empirical data to show that when individuals reciprocate among kin, a spurious effect of kinship is more likely to be detected than a true effect of reciprocity. Further evidence for this bias comes from studies of cooperation in chimpanzees and other primates, which had initially attributed cooperation to kin selection. This conclusion was drawn from the observation that the philopatric sex (females in many Old World cercopithecine monkey and males in chimpanzees) tend to be more cooperative than the dispersing sex (males), which usually interacts more frequently with non-kin [28–30]. However, longer-term observations have instead suggested that while relatedness might facilitate cooperation, it is not determinant in defining social relationships (and is associated cooperative interactions, e.g. grooming [28,31]). These examples do not diminish the importance of kin selection in the evolution of cooperation [32], nor do they necessarily provide support for alternative explanations. However, they highlight the possibility that the observation of kin-biased help might conceal alternative or additional mechanisms, and that their effect may be masked by incomplete data on social interactions.

A key challenge in studying the social drivers of helping behaviour lies in the need for highly detailed social data [33], along with the difficulty of integrating fine-scale social dynamics into longer-term processes (e.g. decisions to help [34]). Until recently, studying social associations in birds was an arduous and largely inefficient task, leading to substantial gaps in knowledge and potentially resulting biases in support for some hypotheses [35], namely kin selection. However, recent advances in tracking animals, such as through GPS-tracking (e.g. [36]), video tracking (e.g. [37]), and the use of radio frequency identification (RFID) technology combined with miniaturized Passive Integrated Transponders (PIT-tags) [38], now allow many individuals—and therefore their relationships—to be tracked simultaneously. These methods, together with appropriate statistical tools (e.g. social network analysis[39,40]), have fundamentally changed the study of social behaviour in birds (and other taxa) and can bring new insights into the study of cooperation in animals [17]. In the specific context of cooperative breeding, these advances can facilitate crucial insights into the social structure and social behaviour during and outside the reproductive season [41–43]. They enable precise quantification of the social context in which helping decisions are made, adding this dimension to the genetic and (relatively large-, or territory-scale) spatial dimensions that have been commonly considered. Integrating these three components —genetic, social, and spatial—provides a more comprehensive understanding of the mechanistic basis of helping (e.g. social bonds, “kin clues” or spatial proximity), while simultaneously providing insights into the ultimate causes of helping (i.e. production of non-descendent kin or increased survival and/or reproduction through direct benefits), and the interplay between the two (see [1,44]). The ability to efficiently collect social association data on cooperative breeding birds could be as revolutionary to properly address hypotheses that rely on the potential benefits of sociality (i.e. “pay-to-stay”, “group augmentation”, “partner choice”, “social prestige”) as the advent of genetic markers were for the study of kin selection.

Hence, in this work we leverage these methodological advances to examine how pre-breeding social ties relate to helping decisions and, in turn, how these decisions relate to social associations after breeding in a colonial and cooperatively breeding bird with kin-biased helping, the sociable weaver *Philetairus socius*. In this species multiple breeding pairs reproduce within a colony and can be assisted by helpers. We deployed PIT tags and feeders fitted with RFID loggers to collect social associations while foraging, and use social network analysis to quantify the social ties between breeders and other colony members. We first examined how social associations between breeders and their helpers varied across time (before, during and after breeding) and how they compared to social associations between breeders and their non-helping kin. We then used a data-driven simulation approach to infer the potential role of prior associations in determining helping decisions while controlling for genetic relatedness and for potential saturation of helping opportunities. Finally, we tested if social associations prior to breeding are related to the amount of helping provided at the nest and if this cooperative investment, in turn, is linked to the social associations between breeders and helpers after chicks have fledged. We predicted that if social associations play a role in driving helping decisions, individuals should help breeders with whom they share the strongest social bonds, and that the pre-breeding social ties between helpers and breeders should be stronger than those between breeders and non-helping kin. We also predicted that if helping brings direct benefits through improved social associations, breeders should stay more strongly connected with their helpers after reproduction (when compared to non-helping kin). Finally, we make similar predictions regarding the amount of help. Individuals should help more according to their association strength with the breeding pair, and changes in social relationship between the periods before and after reproduction should be positively correlated with the amount of help provided.

## Methods

### Study species and population

We worked on a sociable weavers’ population located ca. 6km southeast of Kimberley, in the Northern Cape Province, South Africa. Sociable weavers reproduce and roost throughout the year in large communal nests or “colonies” usually built on *Acacia* (*Vachellia*) *erioloba* trees. In our population, these colonies can host ca. 10-160 individuals, and are composed of multiple independent chambers (up to 78) separated by a few centimetres (ca. 20cm). Adult sociable weavers feed on seeds and insects [45], but provide mainly insects to the nestlings. Both helpers and breeders provide food to the nestlings [46] and help to maintain the nest structure through building and sanitation, defending it from both nest predators and conspecifics [23]. Most helpers are related to one of the breeders, but up to ca. 14% of helpers can be more distant kin or unrelated (R≤0.125 [23]). Sociable weavers from the same colony often forage together, but individuals can also forage alone, in small groups, or in very large flocks containing individuals from more than one colony [47,48].

Regular captures of the study population go back to 1993. Since 2008 captures take place before breeding (August and September) and individuals are ringed with a unique metal ring and colour combination, and a blood sample is collected for sexing and genotyping (see [49,50]). During breeding, all chambers in the study colonies are routinely inspected to identify new clutches, hatching dates, and to record clutches and broods’ fates (fledged, depredated or failed [50]). All chambers are visited when the oldest nestling reaches 9 days old to ring and collect blood samples and again at 17 days-old to colour-ring chicks and record how many reached fledgling stage. Chicks can stay in the chamber until ca. 25 days old. In five (out of 15) colonies in the study population, individuals are additionally fitted with a PIT-tag at day 17 or during the annual captures (if first captured as adults, [47]).

### Social associations’ data collection

To collect data on foraging associations we used artificial feeding stations located ca. 80-205 meters from the colonies [47]. In short, perches with RFID antennas were placed in front of bird feeders in groups of four to form a “feeding box” where birds feed in very close proximity from each other (ca. 5 cm). Feeding stations initially comprised two feeding boxes (between September 2017 and April 2018) and were later modified to include four feeding boxes in order to collect data that was more suitable for larger colonies and the foraging behaviour of the study species (see [47]). We attempted to collect foraging associations every 3 days from September 2017 to December 2019.

We used the detections at these feeding stations to build social networks [40]. The edges of the networks were calculated based on the proportion of time that individuals spent feeding together in the same feeding box given their availability to feed together [47]. This was done by adding the time that two individuals spent feeding simultaneously divided by the sum of time when at least one of these two individuals were present at the feeding boxes (the Simple Ratio Index [51]).

### Nestling provisioning and breeding group determination

During the nestling period, we placed video cameras (Sony HandyCam®) underneath each colony, pointed at individual chamber with nestlings to record colour-ringed adult birds entering the chambers to feed them. Individuals were considered to belong to the same breeding group if they were seen visiting the same chamber to feed the nestlings three times or more. This threshold was applied to avoid including prospecting birds or intruders as group members [23]. Chambers were recorded during four or more hours over the course of two or more days (with each recording typically lasting two hours). Recordings were conducted between days 7-19 of the nestling period (day 1 being the hatching of the first egg).

### Breeder identification and relatedness

To distinguish between helpers and breeders and determine the relatedness between adults and chicks we conducted genotyping analyses on a total of 6169 individuals during 10 years as part of a long-term study on this population. Paternity was determined using a combination of genetic markers (16 microsatellite, see [49]), video analyses and individuals’ breeding history (see [23,50,52]), since extra-pair paternity is uncommon in this population (<3%; [53]). Using this large pedigree dataset, we inferred familiar relationships between the individuals visiting the chambers and the chicks and from those relationships we defined three relatedness categories: R=0.5, R=0.25 and R≤0.125.

### Associations over time

We quantified foraging association rates all year-round in order to capture periods of breeding and non-breeding activity. This allowed us to have association rates before, during, and after periods of breeding activity for each combination of birds (hereafter a “dyad”) in the colony. Using these, we could compare patterns of association among helpers and breeders to those of breeders and their kin that were not observed helping them. We used moving time windows encompassing one-month period (30 days) of data collected at the artificial feeding stations and moving this period every 15 days (i.e. two consecutive windows would overlap 15 days of data). This was done to ensure that each window had sufficient data to generate robust associations. This resulted in 52.4±1.6 (mean±SD) networks per colony.

We analysed our social association data by focusing on each breeder individually. For each focal breeder, we extracted all the connections (edges) from the networks in which this breeder was present (e.g. breeder A could have been present in the first 10 networks, encompassing 135 days of colony B, and then disappeared from the population). Then, we calculated the time difference (in days) between the midpoint of each network (i.e. day 15 of the 30 days of the network) and the day the chicks of that breeder hatched. For each edge, we further noted the relatedness (R≤0.125, R=0.25 or R=0.5) between the focal breeder and the non-breeder individual, whether the non-breeder individual was a future helper, a past helper, or whether it never helped that breeder, and sex similarity (i.e. a same sex or mixed sex dyad). We then categorised all the edges according to the time distance between the mid-point date of the network to the hatching day of the chicks of the breeder into 15-day time bins ranging from 150 days prior to hatching to 150 days post-hatching (n=20 bins). This means that edges from different time periods and individuals of different colonies can be grouped in the same time bin.

Based on this data set we analysed three dependent variables representing the breeder helper or non-helper kin strength of social association while foraging:

#### i) Zero-non-zero edges

We first tested if an individual could be a helper of the breeder without ever being found feeding in close proximity at the artificial feeding stations (i.e. having a 0-edge value). Overall, our networks were composed by many zeros (72,31%±15,99% mean±SD of edges had a value of 0) meaning that even though we often observe the whole colony foraging as a single flock, many individuals are never found feeding in very close proximity to each other. For each of the 20 time bins we tested if breeder-helper edges were less likely to be 0 (i.e. never found together) than the breeder-non helper kin (R>0.125) edges by running a generalized linear mixed models with a binomial distribution with edge value (0 or 1) as dependent variable. We included edge category (helper vs. non-helper edge) and sex similarity as fixed effects. As random effects we included colony ID and the time period of the network (as edges on the same time bin can come from different time periods). For this, and all analyses below, we restricted associations to edges between breeders and kin, because edges with non-kin were generally clearly much weaker. Nevertheless, because non-kin associations provide a useful perspective on the expected strength between any two individuals from the same colony, we include them in all our figures in the results section.

Given that edges of a social network are based on non-independent observations, which have inherent uncertainty associated with them [54], we conducted permutation tests to determine the significance of the GLMM’s estimates of the dependent variables. Specifically, we generated for each time bin replicated datasets in which the associations (i.e. 0 or 1) among individuals was randomized (i.e. a node permutation). We considered the effect of the dependent variable to be statistically significant if the observed estimate fell outside the 95% range of the estimates generated by 1000 randomized datasets. Randomizations were constrained to occur only between edges that were extracted from the same network and that had the same sex type (e.g. male-female edges were only swapped with other male-female edges; constrained node permutations).

#### (ii) Preferred partnership

We then tested if helper-breeder associations were particularly strong compared to other edges of the network. In order to do this, we took a “preferred partnerships” approach in which edges of the network that were twice as larger as the network mean were considered to be especially high (e.g. [31,55,56]). We tested if these preferred associations were more common among breeder-helpers than breeder-non helper kin associations by taking a similar approach as explained above, but using preferred association (0 or 1) as dependent variable in the GLMM.

For these two analyses above, we conducted separate analyses for male and female focal individuals (i.e. breeders). For all, we also only considered helpers that were helping a given breeder for the first time. This separation allowed to infer about the timing in which helper-breeder associations might start to form without the confounding effect from potential carry over effects from previous helping associations. Therefore, helpers helping a second time were included in the data to generate the social networks but their edges were not considered for the statistical analyses. Furthermore, as helpers are on average more related to breeders than non-helping kin (relatedness for breeder male-helpers: 0.43±0.13; breeder male-non-helping kin: 0.37±0.13; Wilcoxon test: W = 75087, p<0.001; N=66 helpers and 243 non-helping kin), in order to avoid a potential cofounding effect between social associations and relatedness we repeated the two analyses using only R=0.5 individuals. As the results were similar to the ones obtained with the whole dataset, we included them in the supplementary material (Figure 1S).

**Figure 1.**
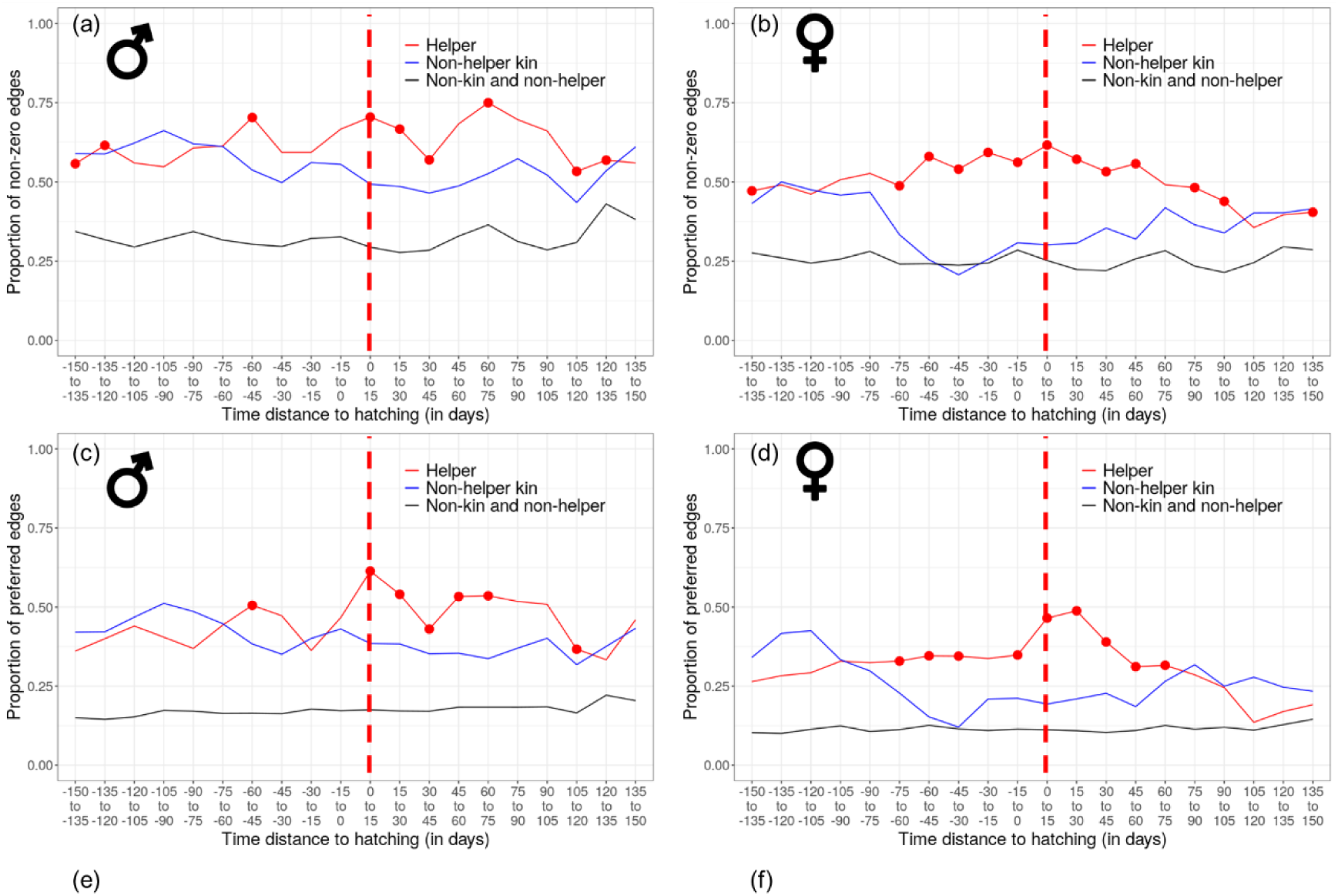
Comparison between helper-breeder, non-helper kin-breeder and non-kin non-helper-breeder edges across time. On the left the results for male breeders and on the right for female breeders. The vertical dashed red line represents the hatching time. The horizontal solid red lines represent the helper-breeder affiliations, analysed in two different ways: proportion of non-zero edges (a and b) and preferred edges (c and d. red lines represent helper-breeder edges (N=95 breeder male edges; N=88 breeder female edges). Blue lines represent non-helping kin-breeder edges (N=243 breeder male edges and N=75 breeder female edges). Black line represents non-kin non-helper edges (N=3941 breeder male and N=3517 breeder female edges), which were not included in the (G)LMMs, but are represented here for illustrative purposes. Red dots indicate time windows in which the edge category helper-breeder was significantly (p<0.05) more likely to be a non-zero edge (a and b) or a preferred edge (c and d) See Figure 2S for time windows in which the opposite was found.

### Data-driven simulation of helping decisions

A simple approach to infer the role of pre-hatching associations in defining helping interactions would be to test if helpers always assisted the breeders with whom they share the strongest social bond (similar to [57], but using social affiliations instead of relatedness). However, such an approach relies on the assumption that helping associations are independent from each other within a colony (or population). This may not be the case if we consider that chicks have a limited amount of food that they can eat and that helpers might be prevented [23] or refrain from helping (e.g. if enough adults are already attending the nestlings; see [58]). Thus, it is prudent to consider that an individual assisting a given breeder reduces the possibilities for other individuals to help that same breeder, making the helping associations non-independent from one-another. In this scenario, the connections between breeders and helpers may not be fully maximized at the individual level—meaning that individuals might not always (be able to) help their closest social partner—but these connections can still influence helping relationships. We therefore used a data driven simulation approach to test for a potential role of social associations in driving helping decisions while considering the potential non-independence of helping associations.

Our simulations used time windows moving day by day. We used this time step because helping opportunities change drastically on a daily basis by having new chicks hatching, disappearing and helping decisions made every day. For each time window we selected all the associations collected at the artificial feeding stations in the previous two months and selected all the active breeding chambers in which the chicks hatched within 10 days from the time window date (last day of the two-months period of the social associations data). We built social networks using the two-month data period and filtered the network to only include the edges from the individuals seen feeding the chicks (i.e. helpers and breeders) that hatched within the 10-day period. Then for all possible dyads of helpers of different chambers (i.e. broods) we “swapped” them if, by swapping them, each would have a higher edge value with the “new” breeder compared with the breeder that they were observed helping. We then compared the number of such swaps that we could make with the number of swaps (using the same rule) that are possible in randomised networks (see below). If helping associations are influenced by social associations, we expect to 1) to find a very small number of “swaps” that would result in higher association strengths for both “swapped” helpers, and 2) find a significant smaller number of “swaps” between the number of observed and the distribution of “swaps” from the randomized networks. This is because randomized networks should generate many more possible swaps than if helpers try to help the breeder with whom they have the strongest social association. If we observe fewer “swaps” (on the “real networks”) than the ones obtained from the randomizations, we can conclude that sociable weavers associate with the breeders with the strongest affiliation and, when they do not, it is because there are more helpers which have those breeders as their favourite than the number of individuals that can help that breeder.

For randomisations, we randomly re-allocated individuals within the same sex type (i.e. male-female, male-male or female-female) and with the same relatedness value (R≤0.125, R=0.25 or R=0.5), i.e., a constrained node permutation [54]. Constraining the randomizations to occur only among same relatedness values and sex category controls for the potential confounding effect arising from the positive correlation between social associations and kinship among males (see results on “foraging associations over time”) and the tendency for stronger associations among males. As most helpers are related to the breeders (see [23]) and relationships among kin are stronger than among non-kin (see Results), not controlling for relatedness (and sex) might overestimate the number of “swaps” simply because we would be attributing edge values to breeders-helpers that are generally lower than the ones observed for kin relationships (which include most of the helping associations).

To simplify the simulations we excluded helpers feeding at more than one nest simultaneously (which is uncommon in our population). We also ran the simulations separately for male and female breeders as helping decisions are often hypothesise to be dependent on sex of the breeder that the helpers are relate to. The two-month period for the social associations was chosen based on the results of the previous analyses (described in the “Foraging associations over time”) that give little support for a specifically stronger helper-breeder edges for time periods larger than two months before the hatching date.

### Foraging associations and nestling provisioning intensity

We tested if foraging associations could predict helping investment by testing if helpers’ nestling provisioning rates (i.e. number of times that helpers come to the nest to feed per hour) and nest attendance (i.e. number of different days that helpers were seen feeding the chicks) relate with helper-breeder’s associations prior to hatching of the chicks. In the sociable weaver, helpers progressively join the breeding activities and might not be recorded feeding the chicks every day [23]. Analysing nest attendance gives additional insight into the commonly used nestling provisioning rates, as individuals might bring a high number of preys per hour to the nest, but might not feed the chicks every day.

For each helper we built a social network using the two-month period of data collection at the artificial feeders prior to the hatching date of a given nest, and the two-month period after the chicks fledged or died. This allowed us to determine the edge strength between all helpers and breeders before and after the nestling period. We tested if breeder-helper edge strength was related with nestling provisioning rate and nest attendance using linear mixed models and generalized linear mixed models with a binomial distribution, respectively.

For the nestling provisioning rate, we used as dependent variable the number of feeding visits (log-transformed) by the helper in the two hours of video and as independent variable edge strength to the breeder (before or after the nestling period). There were no zero values for nestling provisioning because here we are only determining how much effort helpers put into feeding nestlings when helpers show up to provide to them (which is complementary to the nest attendance analyses below). For this and the nest attendance analyses below, we included nestling provisioning as dependent variable instead of edge strength because nestling provisioning can be dependent on many factors that need to be accounted for including: sex of the helper, age of the helper, maximum daily temperature, group size, the number of chicks in the nest and the chicks’ age (in days), which were all also included as independent variables. We also included colony ID, helper ID and breeding period (defined as peaks of breeding activity; see [23]) as random effects. In addition to the before and after the nestling period models, we ran a third model using the difference between the helper-breeder edge strength before and the helper-breeder edge strength after the chicks have fledged or died. The aim was to test if individuals that provide at higher rates would potentially induce a larger change in the association strength with the breeders.

For nest attendance we used a generalized linear mixed model with a binomial distribution and a binary outcome with 1 for appearing feeding the chicks every day that the nest was recorded and 0 for missing at least one day of provisioning as dependent variable. We included edge strength (before or after) or difference in edge strength (in three different models), the number of days that the nest was recorded (as the more a nest is recorded the more likely it is for one adult to be absent on a single day), the age of the chicks in which the first recording was made (as many helpers only start to feed in the later stages of the nestling period), sex and age of the helper. We included as random effects colony ID, helper ID and breeding period as independent variables.

We ran different models for breeder male and breeder female and determined the significance of the variable of interest (edge strength before or after and difference in edge strength) using randomizations as explained in “Foraging associations over time” section, but constraining the randomizations to occur only between individuals of the same sex and that helped in the same breeding period.

## Results

### Foraging associations over time

*(i) Zero-non Zero edges.* Helper-female breeder edges were more likely to be non-zero edges both before (from –75 days before hatching) and after hatching (up to 105 days; Figure 1b) than non-helper-female breeder edges (among kin). This means that helpers are more likely to have associated with the breeding female at least once before, during and after reproduction compared to individuals that are genetically related to the female breeder but did not help. On the other hand, for helper-male breeder edges, this effect was only significantly detected from the beginning of the nestling period (hatching day) until 75 days after hatching (Figure 1a; Figure 2S). Prior to hatching, for male breeders, although there was a positive trend, this effect was not significant (Figure 1a and Figure 2S), giving less support for helpers to be significantly more associated with breeder males before the nestling period when compared with non-helping kin. Associations between breeders and non-helper-non-kin clearly had a higher proportion of zero edges than breeders with helpers and non-helper kin (Figure 2a, although we did not include these edges in the GLMMs). When considering all the time periods before hatching, 19.4% (21 out of 108) and 19.8% (19 out of 96) of all breeder male-helper and breeder female-helper edges, respectively, were always zero. This suggests that nearly 20% of helpers can assist the chicks of the breeder without any detectable social links at the feeders with the breeders, when considering the associations separately for each breeder. Nevertheless, this value decreases to only 4,1% (3 out of 70) when considering the connections to either one of the breeder male or breeder female from the same nest, suggesting that helpers are almost always an individual that was seen foraging with one or both breeders before the chicks hatched. By contrast, when considering breeder-non helper kin edges, 26.9% (60 out of 223) for the breeder male and 52.8% (37 out of 70) for the breeder female were always zero. When considering all non-helper kin of either the male or the female breeder, 39% (67 out of 168) were always a zero edge with either the breeder male or the breeder female of the same nest. This result indicates that non-helper kin are more likely to be never seen feeding in close proximity with the breeders than helper kin.
*(ii) Preferred partnerships approach.* Similar results were found with this approach. Helper-breeder males were more likely to be preferred edges than breeder-non helper kin, but only after hatching (up to 75 days; although there was a positive trend for before hatching; Figure 1c). For breeder females, this interval was larger, encompassing the 75 days before hatching up to 75 days after (Figure 1d). When analysing all the time periods before hatching, 77.8% (84 out of 108) and 74.0% (71 out of 96) of all breeder-helper edges were at some point a preferred edge of the male breeder and female breeder, respectively. When considering both helper-male breeder or helper-female breeder of the same chamber, 95.6% (70 out of 73) were a preferred edge. This means that helpers were almost always at some point a previously preferred edge of the breeder male or the breeder female or both. By contrast, 63.7% (142 out of 223) and 40.0% (28 out of 70) of non-helper kin were previously a preferred edge of the breeding male and the breeding female, respectively. When considering either breeder male or breeder female edges of the same chamber only 50.0% (84 out of 168) were a previously preferred edge.

### Data driven simulation of helping decisions

In our simulations there were a total 137 potential swaps that could be done when considering helper-breeder male edges and 190 for helper-breeder female edges. From those potential swaps, 4 (2.9%) and 7 (3.7%) were possible to perform (for male and female breeders, respectively), i.e. by swapping these helpers, both helpers would have a stronger connection with the breeders they would attend compared to the observed connection. For both male and female breeders, this value was much smaller than the ones obtained from randomized networks controlling for sex and relatedness (p≤0.001; Figure 2a and b). Even without considering the potential non-independence of helping associations, a simpler analysis—focusing on whether helpers assist the breeder with whom they share the strongest bond—still reveals a similar pattern. The majority of helpers assist the breeder they are more socially connected to. Specifically, 66 out of 98 (67.3%) and 91 out of 125 (72.8%) helpers were seen assisting the breeder male and breeder female, respectively, with whom they shared the strongest bond. Together, these results suggest that weavers tend to help the breeders they are more strongly connected to, unless other potential helpers also have a strong connection with those same breeders.

**Figure 2.**
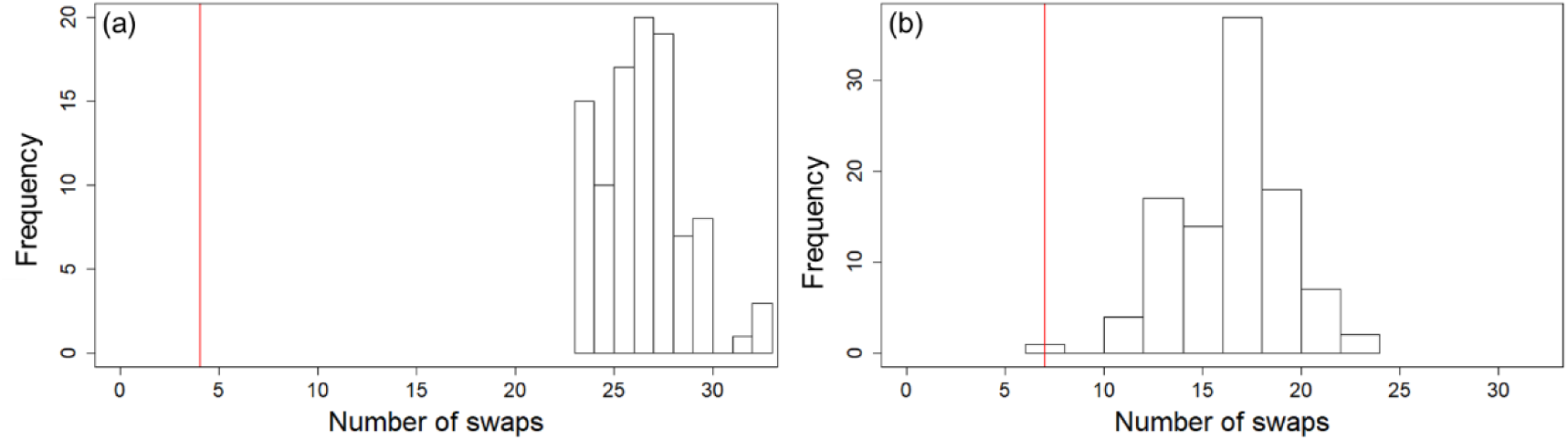
Distribution of the number of pairs of helpers that could be “swapped” based on association strength (red line) and when accounting only for sex and kinship (histograms). Results are shown for (a) breeder males and (b) breeder females.

### Foraging associations and nestling provisioning

Regarding nestling provisioning rates, the associations between helpers and breeder males before the hatching day was not significantly related with the amount of food brought to the nest per hour by the helpers (estimate=-2.160, p=0.709; N=138 helpers; Figure 3a). Nevertheless, helpers that fed at higher rates were significantly more associated with the breeder male in the two months after the chicks have fledged (estimate=7.33 p=0.002; N=120; Figure 3c). Furthermore, individuals that fed at higher rates had a greater increase in their association with the breeder male from pre-hatching to post-fledging (estimate= 8.813; p=0.008; N=115; Figure 3e). For female breeders this later effect was not found (estimate=0.695; p=0.78; N=110; Figure 3f), instead helpers that were more associated with the breeding female both before hatching and after fledging fed the chicks at higher rates (before hatching: estimate=11.047; p=0.012; N=128; after hatching: estimate=3.796; p=0.031; N=117; Figure 3b and d).

**Figure 3.**
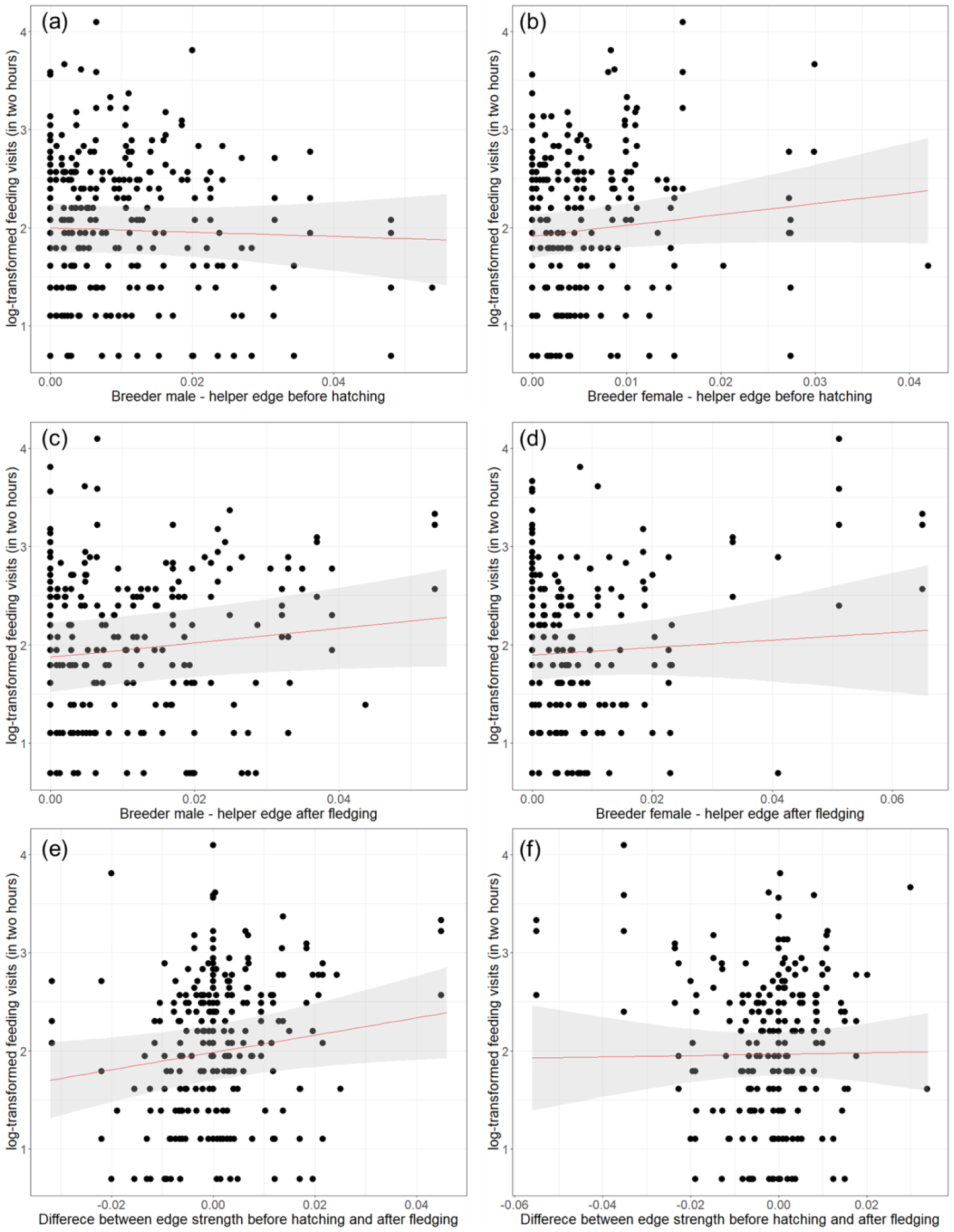
Effect of breeder male-helper (both sexes) (a,c,e) or breeder female-helper (both sexes) (b,d,f) edge strength on feeding visits of the helpers. Edge strength resulting from associations collected two months before hatching (a,b), collected two months after fledging (c,d) and the difference in edge strength before and after fledging (e,f). Black dots represent the raw data, the red line represents the model’s predicted values and the shaded grey area the 95% confidence intervals.

For the attendance analyses, the probability of appearing in all days of recording was not significantly related with the helper-breeder edges before or after hatching or with the difference between both edge types for both male and female breeders (p>0.062 for all analyses; N=138 helpers for male breeder analyses and N=128 helpers for female breeder for before analyses; N=120 and N=117 for after fledging; N=115; N=110 for the difference between before hatching post-fledgling analyses).

## Discussion

In this work we first looked at the associations over time between breeders and either their helpers or their non-helping kin and found that helpers tend to be more strongly associated with the breeding pair both before and after fledging than non-helping kin. Our data-driven simulations further indicated that individuals tend to help the breeders with whom they share the strongest social tie in the colony. We also show that investment in helping (i.e. the number of provisioning visits) is related with the social associations between helpers and breeders. Together these results suggest that social affiliations have a major role in determining helping associations, revealing a link between the strength of social associations and the quality of help given/received and, finally, indicating that social affiliations themselves might be affected by helping interactions.

Breeders were generally more likely to be seen foraging in close proximity at least once with their helpers during the nestling period and after fledging, when compared to their non-helping kin. Similar results were found when considering preferred edges of breeders. Previous work on other species has shown that the quality and type of the social bonds affect the fitness of individuals [11–13]. Nevertheless, studies using social network analysis on cooperative breeding birds remain scarce (but see [42,43,59]), and none have studied the interplay between social associations, kinship and helping at the nest. If helping strengthens social associations, it would provide an avenue by which non-breeding individuals could gain direct fitness.

In the sociable weaver a likely mechanism linking helping, sociality and fitness is communal roosting [60,61]. In this species, birds roost throughout the year in the same chambers used for reproduction, and the size of the roosting group has been shown to have thermoregulatory consequences [62]. Breeding group membership predicts roosting associations [63], suggesting that, by helping, individuals could increase their chances of having access to a roosting chamber (as predicted by “pay-to-stay” hypothesis) or increase their future roosting group size (group augmentation hypothesis). Another mechanism by which helpers could gain direct benefits is if by helping they increase their chances of being chosen as social partners (“partner choice” hypothesis).

Sociable weaver forage in groups and therefore having stronger social ties could facilitate access to food through nepotism [64], harassment avoidance [65], or by obtaining information on new food sources [66,67]. This is especially important for helpers that improve the social ties with the breeding male given that they dominate access to food sources in this species [52]. Other potentially important mechanisms are nepotistic predator protection (e.g. [68,69]), facilitating access to breeding opportunities (e.g. [70]). Here we provide initial evidence that social associations are indeed associated with helping, the next step in future work would be to investigate whether the strengthen of these social ties results in higher survival and/or reproductive success for helpers and/or breeders, thereby identifying the mechanisms underlying those potential fitness benefits.

Our data driven simulations showed that helpers-breeders edges are very close to being maximized at the colony level, meaning that individuals appear to help the (possibly available) breeder with whom they share the strongest social tie. Data driven simulations are limited in their interpretation and can only serve to explore potential mechanisms underlying the observed patterns. This means that alternative mechanisms could also explain this pattern. Nevertheless, these results suggest that social associations provide a potential meaningful context to understand helping decisions. The importance of social context is further supported by another study in this species showing that aggression can explain helping decisions better than kinship [23]. Other anecdotal observations of aggressive interactions between breeders and potential helpers were also reported in other cooperatively breeding species (e.g. [57,71–73]). Taken together, these studies and the results presented in the present study indicate that helping does not occur in a “social vacuum”. These findings call for future experiments on helping decision that take into account the social organization of the species.

The current lack of studies explicitly considering the influence of the social environment on helping in birds is likely to be due to the challenges of studying sociality away from the nest [35]. However, current technologies now enable such studies providing opportunities to investigate how hitherto cryptic social associations influence cooperation. In addition, kin selection, does not predict that breeders might reject help or that there might be competition among individuals to help. Nevertheless, being part of a group is generally predicted to be beneficial for individuals, but the inclusion of additional group members can be costly, ultimately leading to competition for helping or representing added risks to the breeders (e.g. parasitism, cannibalism, infanticide, attract predators). In addition, competition for helping and rejection of help from parents are central to hypotheses such as partner choice and pay-to-stay [14,74,75].

Numerous studies have previously explored the relationship between helping investment and genetic relatedness with mixed results (e.g. [76–88]). The fact that a link between kinship and helping is not always found could be that these studies did not take into account the social context in which helping occurs. In the sociable weaver, kinship correlates with social associations, but does not appear to play a major role in determining the amount of food provided to the chicks [23] and therefore alternative explanations are required to explain variation in helping effort. Here we found that, contrary to expectations, social associations were not related with the attendance of helpers (i.e. the probability of appearing all days to feed the chicks), but nestling provisioning rates by helpers (i.e. the investment when helping) were positively correlated with the social association with the breeders. However, this effect differed according to the sex of the breeder. For breeder females both the associations prior to hatching and post-fledgling were positively related with nestling provisioning rates of the helpers. On the other hand, for breeder males the associations prior to hatching did not predict helping investment, but the post-fledgling associations and the differences between associations post-fledging and prior to hatching were positively correlated with the investment of the helpers. These differences in these patterns of association could be the result of the differences in the strength and number of connections between males and females, as males are philopatric, have more kin and more long-term social bounds than females. However, and regardless of this sex difference, the results from both sexes reveal an association between helping effort and the social bond between helpers and breeders.

In conclusion, our findings suggest that pre-breeding social associations can potentially affect helping decisions and these, in turn, can further influence social associations after reproduction. These results are especially relevant in light of previous studies suggesting that kin selection explains helping behaviour based on the widespread observation that helping is kin-biased. Our findings show that this observation may be concealing potentially relevant direct fitness benefits that individuals can obtain through their social ties.

## Acknowledgements

We would like to thank: Annie Basson, Ryan Olinger, Samuel Perret, Annick Lucas, Cecile Vansteenberghe, Sandra Esteves, Rita Leal, Zoe Tarren and several other field managers, assistants and volunteers for helping with data collection in the field. We further thank Franck Theron for database management. De Beers Consolidated Mines provided access to the Benfontein Reserve. This study was supported by funding from ERC (EU, Consolidator grant 866489), FCT (Portugal, grants IF/01411/2014/CP1256/CT0007 and PTDC/BIA-EVF/5249/2014) and DST-NRF Centre of Excellence at the Fitzpatrick Institute of African Ornithology University of Cape Town to RC, ANR (France, grant 19-CE02-0014-01) to CD. This project is part of the OSU-OREME, and long-term Studies in Ecology and Evolution (SEE-Life) program of the CNRS. RC, RF and ACF were funded by FCT (CEECIND/03451/2018, SFRH/BD/130134/2017 and SFRH/BD/122106/2016, respectively).

## Authors’ contributions

ACF, CD, RC conceived the original idea. ACF, CD, RC and DRF contributed to the methodological design for the social network data. LRS analysed most of the video data for nestling provisioning. ACF, CD, RC, LRS and RF contributed to the long-term data for the pedigree determination. ACF, LRS and RF collected the field data. CD and RC managed the long-term field project that provided the data for this work. ACF run the statistical analyses with the contribution of CD, RC and DRF. ACF led the writing of the first version of the manuscript with the input from CD, RC and DRF. All authors further contributed to improve the subsequent versions of the manuscript.

## Ethical note

Sociable weavers used in this work were captured using standardized protocols approved by the Northern Cape Nature Conservation (permit FAUNA 1638/2015, 0825/2016, 0212/2017, 0684/2019 and 0059/2021) and the Ethics Committee of the University of Cape Town (permit 181 2014/V1/RC and 2018/V20/RC).

