## Supplementary Material for "Foraging associations are related with helping interactions in a cooperatively breeding bird"

[c]CIBIO, Centro de Investigação em Biodiversidade e Recursos Genéticos, InBIO Laboratório Associado, Campus de Vairão, Universidade do Porto, 4485-661 Vairão, Portugal

[d]BIOPOLIS Program in Genomics, Biodiversity and Land Planning, CIBIO, Campus de Vairão, 4485-661 Vairão, Portugal

[e]Department of Collective Behaviour, Max Planck Institute of Animal Behavior, Konstanz Am Obstberg 1, 78315 Radolfzell, Germany

[f]Division of Ecology and Evolution, Research School of Biology, Australian National University, 46 Sullivans Creek Road, Canberra ACT 2600, Australia

[g]Department of Biology, Norwegian University of Science and Technology, Trondheim, Norway

[h]FitzPatrick Institute of African Ornithology, DST-NRF Centre of Excellence, University of Cape Town, Rondebosch 7701, South Africa

### Supplementary material

Foraging associations over time considering only R=0.5 individuals

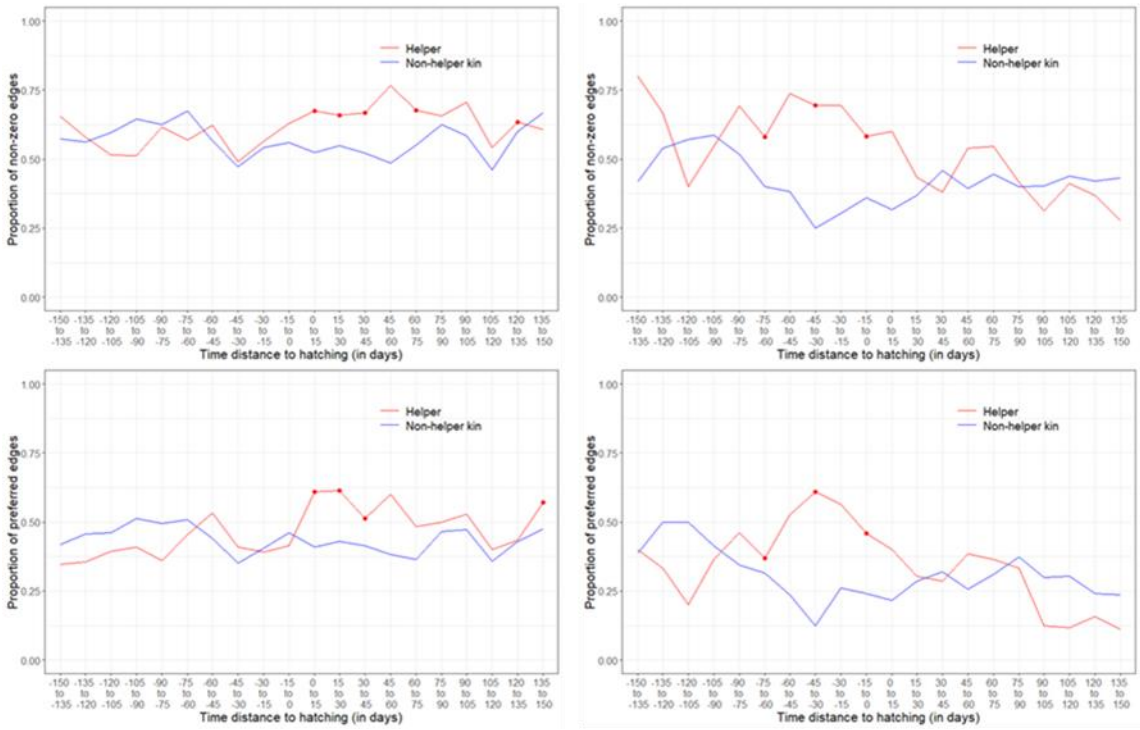


**Figure 1S.** Comparison between helper-breeder and non-helper kin-breeder edges across time, only for dyads of R=0.5. On the left the results for male breeders and on the right for female breeders. The red lines represent the helper-breeder affiliations, analysed in three different ways: proportion of non-zero edges (a and b), preferred edges (c and d). Red lines represent helper-breeder edges (N=51 breeder male edges; N=24 breeder female edges). Blue lines represent non-helping kin-breeder edges (N=116 breeder male edges and N=48 breeder female edges). Red dots indicate time windows in which the edge category helper-breeder was significantly ($p<0.05$) more likely to be a non-zero edge (a and b) or a preferred edge (c and d). In comparison with Figure 1 (i.e. containing the all edges and not only R=0.5 edges) the results are similar, but there are less time bins that are significantly different between breeder-helper and non-helper kin-breeder when considering only R=0.5 edges. Nevertheless, this can be a consequence of a drastic reduction in the sample size, as for the time bins that are no longer significant, they generally still follow the same trend as in Figure 1.

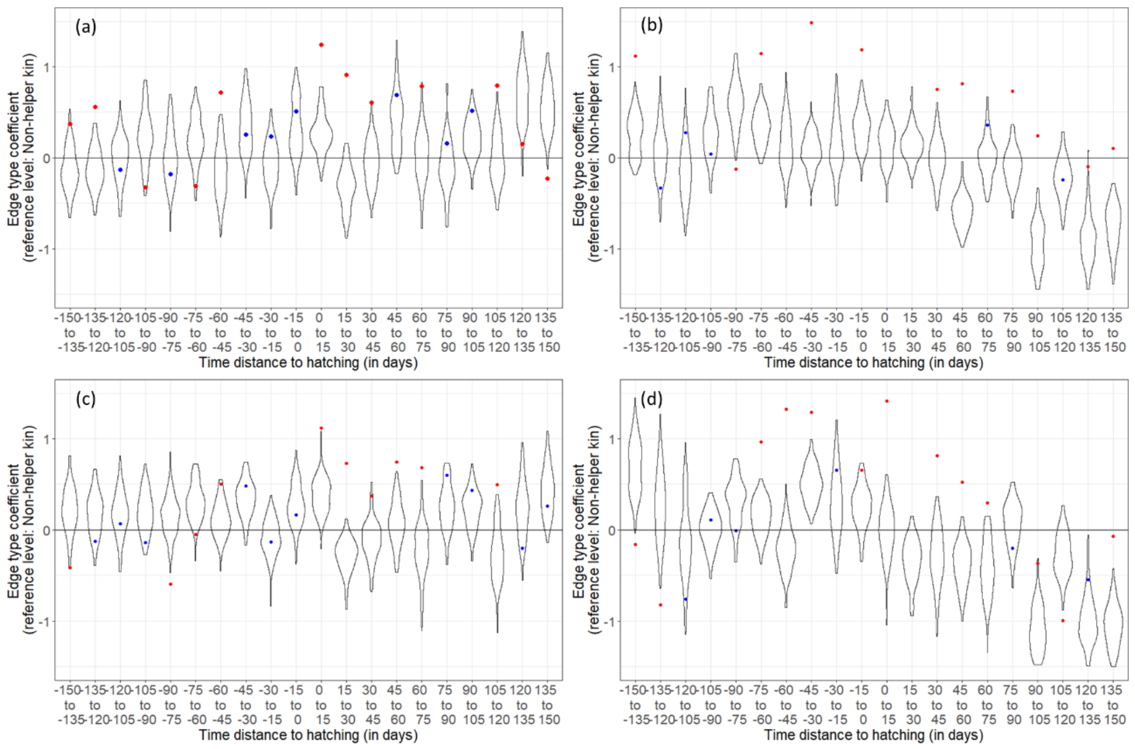


**Figure 2S**. Observed estimates (dots) and respective estimates obtained from 100 randomized networks for each time window (violins) for the variable edge type (helper vs non-helping kin) in the (G)LMMs used in the foraging associations over time analyses. Red dots indicate significance ($p<0.05$) and blue dots represent non-significant estimates. On the left side results are shown for helper-breeder male edges and on the right side for helper-breeder female edges. (a) and (b) Probability of being a non-zero edge. (c) and (d) Probability of being a preferred edge.
